# A NEW CASE OF DIPLOIDY WITHIN A HAPLOID GENUS OF ENTOMOPATHOGENIC FUNGI

**DOI:** 10.1101/2021.03.15.435421

**Authors:** Knud Nor Nielsen, João Felipe Moreira Salgado, Myrsini Eirini Natsopoulou, Jason E Stajich, Henrik H. De Fine Licht

## Abstract

Fungi in the genus *Metarhizium* are soil-borne plant-root endophytes and rhizosphere colonisers, but also potent insect pathogens with highly variable host ranges. These ascomycete fungi are predominantly asexually reproducing and ancestrally haploid, but two independent origins of persistent diploidy within the Coleoptera-infecting *M. majus* species complex are known and has been attributed to incomplete chromosomal segregation following meiosis during the sexual cycle. There is also evidence for infrequent sexual cycles in the locust-specific pathogenic fungus *Metarhizium acridum* (Hypocreales: Clavicipitaceae), which is an important entomopathogenic biocontrol agent used for the control of grasshoppers in agricultural systems as an alternative to chemical control. Here, we show that the genome of the *M. acridum* isolate ARSEF 324, which is formulated and commercially utilised under the name ‘Green Guard’, is functionally diploid. We used single-molecule real-time (SMRT) sequencing technology to complete a high-quality assembly of ARSEF 324. Kmer frequencies, intragenomic collinearity between contigs and single nucleotide variant read depths across the genome revealed the first incidence of diploidy described within the species *M. acridum*. The haploid assembly of 44.7 Mb consisting of 20.8% repetitive elements, which is the highest proportion described of any *Metarhizium* species. The genome assembly and the inferred diploid state, can shed light on past research on this strain and could fuel future investigation into the fitness landscape of aberrant ploidy levels, not least in the contest of biocontrol agents.

## INTRODUCTION

The duplication of single genes, genomic segments, chromosomes, and whole-genome duplications (WGD) has played a vital role in eukaryotic evolution by providing genetic material for adaptation. Throughout the evolution, recurrent WGD events have been linked to the emergence, diversification and survival of species (Crow and Wagner, 2006). For example, WGD has been coupled with the emergence of angiosperms within plants (De Bodt, Maere and Van de Peer, 2005), and with the emergence of vertebrates, gnathostomes and teleosts within metazoan chordate evolution (Sacerdot *et al.*, 2018).

A well-described ancient WGD event in the fungal kingdom is the ancient diploidisation that lead to the formation of the *Saccharomyces* genus (Wolfe and Shields, 1997; Dietrich, 2004; Kellis, Birren and Lander, 2004). While some WGD events lead to stable polyploids, as in the case of *Saccharomyces* where an ancient hybridization event is thought to have provided stability and fertility to the new diploid (Marcet-houben and Gabaldón, 2015), duplications are usually perceived as transient. Duplicated genes enter into a cycle with subsequent reductions by loss or mutational decay leading to pseudogenisation or neofunctionalisation (Lynch, 2000; Levasseur and Pontarotti, 2011). It should be noted that it can be difficult to distinguish ancient WGD events from cases of multiple segmental duplications caused by transposon activity (Roelofs *et al.*, 2020). Regardless of origin, duplications have long been recognised as a driver of evolution by providing the material for natural selection to work on (Bridges, 1919; Metz, 1947; Ohno, 1970; Crow and Wagner, 2006).

WGD arise from abnormal cell cycles post chromosome duplication, either by the absence of mitotic division or cytokinesis. The duplication itself can occur by auto- or allopolyploidisation. The latter involves the fusion of individual cells and subsequent fusion of nuclei bringing together the variants accumulated between the genetically distinct individuals. In contrast, autopolyploidisation, or endoreplication, only involves a single cell and has been associated with environmental stress. Within plants, several studies have shown that endoreplication can be induced by abiotic stress, such as heat (Monjardino, Smith and Jones, 2006), drought (Cookson, Radziejwoski and Granier, 2006), elevated salinity (Barkla *et al.*, 2018), and toxins (Biskup and Izmaiłow, 2004). The duplication is thought to mediate an acute increase in the transcription of metabolic and stress-mitigating genes (Scholes and Paige, 2015). The human pathogenic fungi *Candida albicans* and *Cryptococcus neoformans* have been shown to repeatedly gain resistance to antifungal drugs such as fluconazole through chromosomal duplications (Sionov *et al.*, 2010; Kronstad *et al.*, 2011). *C. neoformans* can also evade phagocytosis within the lungs by creating titan cells (Okagaki *et al.*, 2010; Zaragoza *et al.*, 2010). The increased cell size is induced by multiple rounds of endoreplication and is observed in up to one-fifth of the *C. neoformans* cells residing in infected lung tissue (Okagaki *et al.*, 2010). Transient somatic endoreduplication has also been recognised as a driver of developmental change of some cell types in plants, insects and mammals (Fox and Duronio, 2013). Examples of somatic endoreplication include the giant cells of the mammalian trophoblast that develops into a significant part of the placenta, and the >1000x sister chromatids observed in the salivary glands of *Drosophila melanogaster* (Sher *et al.*, 2013).

Within fungi, increasing evidence suggests widespread cryptic population-level ploidy variations. Whole-genome sequencing of an extensive collection of 794 natural (wild) isolates of *Saccharomyces cerevisiae* found alternative ploidy levels within 13%, of the otherwise predominately diploid yeast (Peter *et al.*, 2018). The same study found chromosomal copy number variation (CCNV or aneuploidy) in 19% of the isolates (193 of the 1,011), reiterating the importance of variation in chromosome numbers for organismal evolution (Mayrose and Lysak, 2020). Although at least some fungi readily undergo endoreplication such levels of genome plasticity is not without constraints. A long-term evolutionary experiment with *S. cerevisiae* showed a convergence from alternative ploidies to the ancestral diploid form (Gerstein *et al.*, 2006). The chytrid fungus *Batrachochytrium dendrobatidis* threatening amphibians worldwide has a high rate of CCNV in nature (Rosenblum *et al.*, 2013). This plasticity has been linked to pathogenicity through lab experiments which showed a gain of chromosomal copies upon induced stress by a characteristic host antimicrobial peptide (Farrer *et al.*, 2013).

While the majority of known fungi are dikaryotic at some stage, diploidy is the exception. Filamentous ascomycetes (Pezizomycotina) are only dikaryotic in specific life stages, namely the ascogenous hyphae and ascocarp, while the bulk of the mycelium remains monokaryotic, and in the majority of species, haploid. Exceptions to haploidy within this important group of fungi are found in the *Phyllactinia*, *Stephensia*, *Xylaria*, *Botrytis* and *Zygosaccharomyces* genera (Albertin and Marullo, 2012), and within the genus *Metarhizium* (Kepler *et al.*, 2016). When these fungi contain two different idiomorphs of the mating-type genes within the same diploid, it indicates mating events that failed to complete meiosis, leading to the observed allopolyploidisation. Diploidy may also arise from the parasexual cycle of fungi, which allows for nuclei fusion (karyogamy) following anastomosis of strains from somatically compatible groups. Contrary to the sexual cycle, fused nuclei of the parasexual cycle do not undergo meiosis, but continue to divide mitotically. The original ploidy level is restored by random chromosome loss, through a series of aneuploidy intermediates (Moore, Robson and Trinci, 2011), which is known from the ‘asexual’ human pathogen *Candida albicans*. Reversion is a gradual process with generations of aneuploidy. Sexual recombination requires different mating-types, whereas the parasexual cycle normally requires compatible identities. The reproductive isolation of the parasexual cycle is enforced by heterokaryon incompatibility protein (HET) domains that abort incompatible hyphal fusion attempts (Glass and Dementhon, 2006). This entails that any parasexual fusion of nuclei will be of closely related and highly similar genomes. Infected insects can be hot spots of fungal parasexuality, and the formation of diploid conidia (Riba *et al.*, 1980; Leal-Bertioli *et al.*, 2000; Wang *et al.*, 2011). Investigating the conidida harvested from an insect coinfected with two fluorescently labelled isolates of the same *M. robertsii* strain revealed that 24% of the conidia investigated were diploid (Wang *et al.*, 2011). Similar observations have been reported from experiments with *M. anisopliae* and *M. majus* (Riba *et al.*, 1980; Leal-Bertioli *et al.*, 2000).

The entomopathogenic fungus *M. acridum* (Hypocreales: Clavicipitaceae) is a specialist with a narrow host range of insects of the order Orthoptera. Host specificity is enforced by the requirement of Orthopteran cuticle signals facilitating spore germination and appressorial formation (Wang and St. Leger, 2005). The *M. acridum* strain ARSEF 324 is the active ingredient in the biocontrol agent ‘Green Guard’ targeting grasshoppers (Acrididae), which is commercially deployed in Africa, Australia and China where these insects are agricultural pests of concern. Within the genus *Metarhizium, M. acridum*, as well as the specialist *M. album*, are considered to be cryptic sexual species (Hu *et al.*, 2014; St. Leger and Wang, 2020). No teleomorphs are known, but genomic analyses have found signatures of an active repeat-induced point mutations (RIP) machinery, an indirect sign of meiosis and sexual recombination (Galagan and Selker, 2004). This is in contrast to the co-generic generalists with wide host ranges, *M. pingshaense*, *M. anisopliae*, *M. robertsii* and *M. brunneum* (the PARB clade, sensu Bischoff et al. 2009), where no RIP footprint is present, consequently causing these species to be considered primarily asexual.

The ascomycete genus *Metarhizium* is generally haploid, but Kepler *et al*. (2014) found two independent origins of stable diploid clades interspersed among lineages comprised entirely of haploid individuals within the larger *Metarhizium guizhouense/majus/taii* clade (MGT clade; sensu Bischoff et al., 2009). All isolates within these diploid taxa were consistently heterozygotic for the analysed microsatellite markers and contained two different ideomorphs of the mating-type genes (MAT1 and MAT2). The presence of both mating-types was interpreted as a mating event that failed to complete meiosis, which lead to the observed allopolyploidisation. The locust-specific *M. acridum* has so far been regarded as entirely haploid, and the only previously available genome for this species was from the haploid strain CQMa 102. This strain was sequenced and assembled into 241 scaffolds (>1 kb; N50, 329.5 kb) containing 1,609 contigs with a total genome size of 38.0 Mb (Gao *et al.*, 2011).

In this study, we describe another instance of diploidy within the genus *Metarhizium* and the first within *M. acridum*. We present a de-duplicated haploid genome assembly of *M. acridum* strain ARSEF 324, which consist of 35 contigs representing a haploid genome of 44.7 Mb. Ploidy is established through K-mer analysis on unassembled sequencing reads, and intragenomic collinearity analysis by all-vs-all mapping of contigs. Euploidy was established through sequencing read depth comparison across each contig, and through estimation of heterozygosity ratios of single nucleotide variants (SNV), both in 10 Kb non-overlapping sliding windows. Finally, using DAPI fluorescence staining we demonstrate that conidia are uninucleate.

## MATERIALS AND METHODS

### Strain and culturing

The *Metarhizium acridum* strain ARSEF 324 was isolated in 1979 from a spur-throated locust (*Austracris guttulosa*) in Australia and obtained from the ARSEF collection (ARS Collection of Entomopathogenic Fungal Cultures, Ithaca, New York). A single-spore isolate was obtained by plating serial-dilutions of conidia from the original culture. Conidia from the single-spore culture were grown in liquid sabouraud dextrose broth with yeast extract (40 g/L dextrose, 10 g/L peptone, 10 g/L yeast extract, pH = 6.5) while being stirred at 170 rpm on a shaking table for 24 hours before DNA extraction. Fluorescence staining with DAPI (blue, 4’,6-diamidino-2-phenylindole) was used to visualise the number of nuclei within mature conidia, as described in (Kepler *et al.*, 2016). Imaging was done on a Zeiss Axioscope microscope (x100 objective) with an AxioCam ICm1, and spores measured using ImageJ v1.53e (Schneider, Rasband and Eliceiri, 2012).

### DNA and RNA extraction and sequencing

Liquid culture filtrate was ground with liquid nitrogen in a mortar and ca. 500 mg finely ground powder transferred to 50 mL Falcon tube. DNA was extracted using a CTAB method by adding 17.5 mL CTAB buffer with 0.1 % 2-Mercaptoethanol and 125 μl Proteinase K and incubated for one hour at 60 °C. After centrifugation for 20 min. at 5.000g at 4 °C, the supernatant was removed and washed with 1 volume of Phenol/chloroform/isoamyl alcohol (25:24:1) followed by two rounds of 1 volume chloroform/isoamyl alcohol (24:1) and centrifuged for 10 min at 11.000 rpm at each step. To remove RNA, 150 μl RNAse was added to the aqueous phase and incubated for 120 min. at 37 °C. To precipitate DNA, 0.6 vol. 2-propanol was added and incubated at −20 °C overnight, before centrifugation at 11.000 rpm at 4 °C for 30 min. The resulting pellet was washed twice with 2 mL 70% Ethanol and centrifuged for 10 min at 11.000 rpm at 4 °C before being suspended in 500 μl TE-buffer. Purity was checked using NanoDrop reading and DNA quantity using a Qubit Broad-Range analysis kit.

A single SMART Cell library was sequenced on a PacBio Sequel platform using the Sequel^®^ Sequencing Kit 2.1 v2 (Sequencing Kit par number 101-309-500). Each continuous long read (CLR) had one passage of sequencing; subsequently, subreads were generated by the removal of adapters and bases were called with the basecaller v5.0.0.6236. Sequencing was performed by Genewiz (Takeley, UK) with a yield from the single SMRT Cell equaling a coverage of 134x, assuming a 45 Mb genome, with a mean subread length of 11.2 Kb. Short read sequence data was obtained using the PCR-free DBNseq™ platform at BGI-tech Copenhagen. Sequencing adapters were removed by the sequencing company, delivering a total of 15,111,206 reads of 150 bp., equaling a coverage of 50x, assuming a 45 Mb genome.

To obtain RNAseq samples to aid in the genome prediction, four replicate flasks with 20 mL media were grown for four days as described above but with 100 rpm agitation. After filtration through a Whatman filter paper (5-13 μm), the fungal material was collected and flash-frozen in liquid nitrogen before being pulverised in a tissue lyzer and extracted using a QIAGEN plant RNeasy kit following manufacturers specifications. The four samples were sequenced separately using the DBNseq™ platform at BGI-tech Copenhagen yielding ca. 16,000,000 paired-end 150 bp reads per sample.

### K-mer analysis and genome assembly

K-mers frequencies within PacBio subread data was established with KMC 3 (Kokot, Długosz and Deorowicz, 2017), and genome size and ploidy were inferred using GenomeScope v2.0 (Ranallo-Benavidez, Jaron and Schatz, 2020). Ploidy was further investigated comparing the sum of kmer pair coverages using Smudgeplot implemented in GenomeScope. The mitochondrion was assembled from 2806 reads mapping to the mitochondrion of *Metarhizium rileyi* strain RCEF 4871 (NCBI assession: NC_047289.1; Zhang et al., 2020). The alignment was done with Minimap2 v2.17(r941) (Li, 2018). Only reads above eight kb, which mapped with more than 70% of their length was kept. The mitochondrion was assembled using Canu v. 2.0 (Koren *et al.*, 2017) with the settings: ‘genomeSize=62k’ ‘corOutCoverage=999’.

Sequence reads were assembled with Canu v. 2.0, using the following parameters to co-assemble haplotypes: genomeSize=45m correctedErrorRate=0.03 corOutCoverage=200 batOptions=-dg 3 -db 3-dr 1 -ca 500 -cp 50. Two rounds of polishing were conducted to improve the assembly, each consisting of mapping raw-reads to the assembly using Minimap2 v.2.6 (Li, 2018), and reducing remaining insertion-deletion and base substitution errors by polishing the consensus sequence using Arrow v2.3.3 (https://github.com/PacificBiosciences/GenomicConsensus). Mirror-reads (putatively an effect of missing adapters during sequencing) were detected in the PacBio subreads set; therefore, reads longer than 40 kb were filtered out prior to assembly. Different levels of the correctedErrorRate parameter were tested (i.e.: 0.020, 0.025, 0.030, 0.035, 0.040). Each of the five assemblies was assessed based on the complementarity of phased contigs. The value 0.030 were chosen for the final assembly (see sup. figure 1).

Contigs were ordered according to length and grouped into primary contigs, forming a haploid representative genome assembly (Haplotig 1 (H1)) and shorter contigs (H2) that mapped to the primary contigs. Primary contigs were identified by an all-versus-all contig mapping on the repeat masked assembly with minimap2 v.2.17. The best match of each contig to a longer contig was assessed from the accumulated length of alignment between each pair of contigs. Primary contigs were manually curated; three contigs were not only subjects to mapping, but also query to other primary contigs. These contigs, and the regions they mapped to had approximately half the read depth of the mean read depth of the other primary contigs. Since the read depth of the subject regions could be raised to the mean read depth, if the query contigs were removed before mapping, these three contigs were assumed to be haplotigs and moved to the H2 group.

### Transcriptome assembly

Adapter and low quality sequences were trimmed from RNA-seq reads with Trimmomatic v.0.36 (Bolger, Lohse and Usadel, 2014) and poly-A sequences were clipped from transcripts using SeqClean [available at http://compbio.dfci.harvard.edu/tgi/software/]. Reads were aligned to the complete assembled genome (H1 + H2) of *M. acridum* ARSEF 324 using HISAT2 (Kim *et al.*, 2019), allowing for a maximum intron length of 3 kb. This was followed by clustering of aligned reads using Trinity v2.8.5 (Grabherr *et al.*, 2011) in a genome-guided *de novo* transcriptomic assembly, using the jaccard clip parameter to reduce transcript fusion. The Trinity assembled transcripts were input to gene prediction training to support genome annotation.

### Genome annotation

A custom repeat library was created by adding *de novo* identified repeats from RepeatModeler (Hubley *et al.*, 2016) to the repeat databases, Dfam_Consensus-20170127 (Hubley *et al.*, 2016) and RepBase-20170127 (Bao, Kojima and Kohany, 2015). RepeatModeler was run on the genome masked by these two public databases, and the iteratively growing custom library, using RepeatMasker v4.0.7 (Smit, Hubley and Green, 2015) with the option [-species] set to fungi. RepeatModeler relies on the three *de novo* repeat finding programs RECON (Bao and Eddy, 2002), RepeatScout (Price, Jones and Pevzner, 2005) and LtrHarvester/Ltr_retriever (Ellinghaus, Kurtz and Willhoeft, 2008). Repeats were identified using RepeatMasker [option: sensitive] and the generated custom repeat library. The custom library was built on the following four *Metarhizium* strains: *M. acridum* CQMa 102, *M. anisopliae* JEF-290, *M. brunneum* ARSEF 4556, *M. rileyi* ARSEF 4871.

Gene prediction and functional annotation of the polished assembly was conducted using the Funannotate pipeline v1.8.4 (https://github.com/nextgenusfs/funannotate). Repeats were identified with RepeatModeler and soft masked using RepeatMasker. Protein evidence from a UniProtKB/Swiss-Prot-curated database (v2021_01) (Bateman *et al.*, 2021) was aligned to the genomes using tBLASTn and Exonerate (Slater and Birney, 2005). Two gene prediction tools were used: AUGUSTUS v3.3.3 (Stanke and Morgenstern, 2005) and GeneMark-ES v4.33 (Besemer and Borodovsky, 2005), with *Fusarium graminearum* as the model for the AUGUSTUS gene predicter and BAKER1 (Hoff *et al.*, 2016) for the training of GeneMark-ES. tRNAs were predicted with tRNAscan-SE (Lowe and Eddy, 1997). Consensus gene models were found with EvidenceModeler (Haas *et al.*, 2008). Gene models were further updated with the ‘update’ step in funannotate which uses the tool PASA (Haas *et al.*, 2008) to further extend untranslated regions and identify alternatively spliced isoforms based on alignments of RNA-Seq assembled transcripts. The completeness of the assembled genome was evaluated through comparison to the 4494 single-copy ortholog genes of the hypocreales_odb10 dataset (Creation date: 2020-08-05, https://busco-data.ezlab.org/v5/data/lineages/), using BUSCO v5 (Simão *et al.*, 2015).

Functional annotations for the predicted proteins were obtained using BLASTP to search the UniProt/SwissProt protein database (v2021_01). Protein families (Pfam) and Gene Ontology (GO) terms were assigned with InterProScan v5.48-83.0 (Jones *et al.*, 2014). Functional predictions were also inferred by alignments to the eggNOG 5 orthology database (Huerta-Cepas *et al.*, 2019) using emapper v2.0.1b-2-g816e190(Huerta-Cepas *et al.*, 2017). The secretome was predicted using SignalP v5.0 (Almagro Armenteros *et al.*, 2019) and Phobius v1.01 (Käll, Krogh and Sonnhammer, 2007) identifying proteins carrying a signal peptide. To further characterise the secretome, putative CAZymes were identified using HMMER v3.3 (Eddy, 2011) and family-specific hidden Markov model (HMM) profiles of dbCAN2 meta server (Lombard *et al.*, 2014). Putative proteases and protease inhibitors were predicted using the MEROPS database (2017-10-04) (Rawlings, Barrett and Finn, 2016). Biosynthetic Gene Clusters (BGCs) were annotated using strict parameters of the antibiotics and Secondary Metabolites Analysis Shell v5 (antiSMASH) (Blin *et al.*, 2019). All functional predictions and annotations were executed through the Funannotate pipeline.

The guanine-cytosine (GC) content was determined in non-overlapping windows across the 35 H1 contigs, using seqkit v0.13.2 (seqkit sliding -s 100 -W 100 | seqkit fx2tab --name –gc). Each window was annotated with the gene or repeat annotation (or no annotation) which it primarily overlapped. This was achieved with bedtools v.2.28.0 option [intersect -wa -wb -f 0.51].

### Variant calling and chromosome copy number variation (CCNV)

The 150 bp PE DBNseq reads were mapped with the Burrow-Wheeler Aligner, BWA-MEM v.0.7.16a (Li, 2013) to the primary haplotig (H1) of the assembled nuclear genome, and PCR duplicates were removed from the bam file using samtools v1.10 (markdup –r) (Li *et al.*, 2009). Coverage and mean depth to reference was assessed using samtools (coverage). Allele frequency distribution across the genome was calculated based on single nucleotide variants called using Bcftools v1.10.2 (Li, 2011a). Variants were called by using a combination of BCFtools’ mpileup’ and ‘call’ using a mapping quality filter of 30, a base quality filter of 20, and a minimum depth of 10, together with default parameters including BAQ (Li, 2011b). Mean SNV allele ratios were summarised in non-overlapping 10 kb windows using a custom R script. Genome annotations and repeat annotations was combined for the mating-type loci including the flanking APN2 and SLA2 genes, for to the genome assemblies of *M. acridum* ARSEF 324 and *M. acridum* CQMa 102 (GenBank accession no. GCA_000187405.1). Synteny between loci was visualised using the R package genoPlotR v0.8.10 (Guy, Roat Kultima and Andersson, 2010).

### Genome comparison

Single copy orthologous genes were identified between H1 and H2, and 11 *Metarhizium* isolates representing eight species, along with two isolates of *Pochonia chlamydosporia* and two isolates used as outgroup, i.e: *Epichloe festucae* strain Fl1, and *Villosiclava virens* strain UV-8b. This was done using OrthoFinder v2.2.7 (Emms and Kelly, 2019). The proteomes of the 16 isolates was obtained from NCBI GenBank; accession numbers are available in the supplementary table 2. Protein sequence alignment was generated with MAFFT v 7.453 as implemented in OrthoFinder. Substitution models for each gene were predicted using ModelFinder (Kalyaanamoorthy et al., 2017) as implemented in IQtree v2.1.2. A subset of 444 known genes with less than 10% gaps within the alignment was concatenated and a maximum likelihood phylogeny with gene specific substitution models using IQtree (Nguyen et al., 2015; Chernomor, von Haeseler and Minh, 2016).

Repeat-Induced point (RIP) mutations indices were calculated using ‘The RIPper’ (http://theripper.hawk.rocks). Three different indices were calculated, based on dinucleotide frequencies: RIP substrate [(CpA+TpG)/(ApC+GpT)]: 0.75 ≥ x, RIP product [TpA / ApT]: x ≥ 1.1 and RIP composite [(TpA/ApT)–((CpA + TpG)/(ApC + GpT))]: x ≥ 0, where x are values indicating RIP on a given sequence. Conservative estimates the RIP affected proportion of the genome are given as regions where all of the above indices indicate RIP (Figure 5).

## RESULTS

### K-MER analysis

The k-mer analysis on the PacBio subreads predicted that the sequenced genome was diploid, and with a haploid length of 45 Mb (Figure 1A). The profile in figure 1A shows two peaks in the frequency of observed unique 21-mers within the sequencing data. The smaller peak, has half the read coverage of the taller peak indicating the presence of heterozygous loci. The genome is estimated to have a heterozygosity rate of 0.5%. Analysing the sum of kmer pair coverages to coverage ratios confirmed the inferred diploidy of the data figure 1B (For full GenomeScope results see supplementary material table 1).

**Fig. 1.**
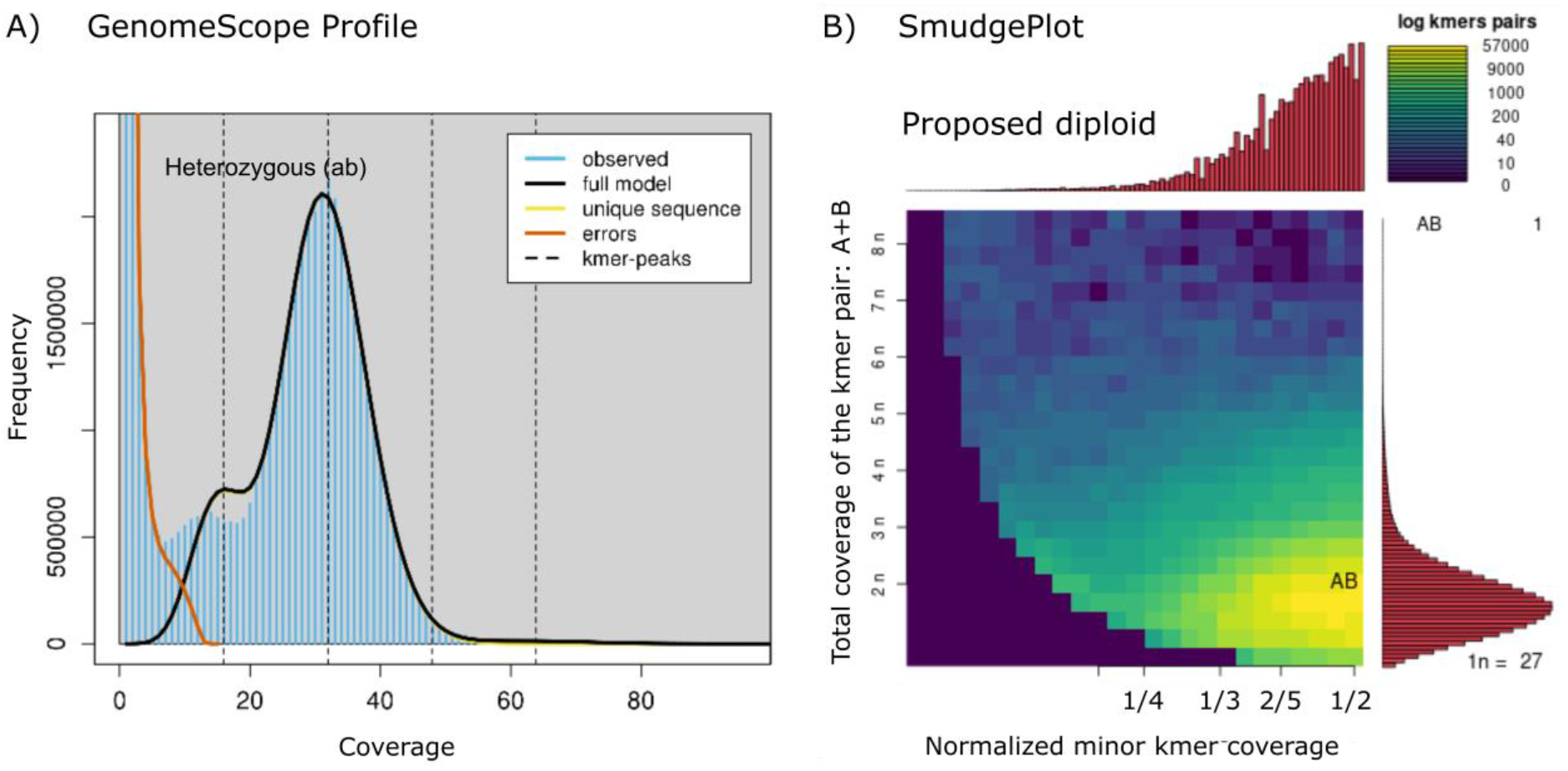
The kmer analysis for Metarhizium acridum ARSEF 324 PacBio subread data revealed a diploid structure of heterozygous kmer pairs. A) Based on the the number of unique 21-mers the genome is estimated to have a haploid length of 45 Mb, with 18.2% repeats. The two peaks of the k-mer frequency profile of the observed data, where the lesser have half the read coverage of the higher peak, indicate heterozygocity, consistent with diploidy. The low frequency of k-mers from heterozygous loci indicate that most 21-mer were heterozygous (0.5%). B) The diploidy of the genome was confermed by Smudgeplot analysis, comparing the sum of kmer pair coverages (CovA+ CovB) to their relative coverage (CovB / (CovA + CovB)). No higher level ploidy was observed. Max and min estimates from the GenomeScope analysis is available in supplementary material table 1.

### Genome assembly

The mitochondrion genome was assembled to a length of 96,363 bp and contained 17 genes. The number and order of genes corresponds with the conserved structure of mito-genomes within Hypocreales (Aguileta *et al.*, 2014). A total of 478,156 filtered Pacbio reads with an average length of 12.7 kb was used to assemble the genome into 35 primary contigs representing a haploid genome of 44.71 Mb. The primary assembly H1 has an L50 of 6, with a N50 of 23.37 Mb, and 4276 complete BUSCO genes, out of the 4494 BUSCO genes sought, equal to 95.1% (Scores in BUSCO format: C:95.1%[S:94.2%,D:0.9%],F:1.4%,M:3.5%,n:4494). Out of the 44.7 Mb in H1, 9.3 Mb (20.8%) was repetitive elements 74.2% of which with a strong GC-bias (<33.2%).

The alternative haplotig (H2) is represented by 565 contigs with a collective size of 39.24 Mb, i.e 88% of the assumed haploid length. This assembly has an L50 of 144, with a N50 of 0.08 Mb. This set of contigs only contains loci sufficiently divergent from H1 to be forked out during the genome assembly. This implies that conserved loci between H1 and H2 will be present in the H1 haplotigs, but largely absent from the H2. This is reflected in the associated BUSCO scores where 43.7% of the BUSCO genes are missing from H2 (C:54.1%[S:51.7%,D:2.4%],F:2.2%,M:43.7%,n:4494). The outer band of the outer tracks of figure 2 depict the size of the 35 primary contigs representing the complete haploid genome. The second band depicts the mapping of the 565 H2 contigs to H1, indicating their redundancy (Figure 2).

**Fig. 2.**
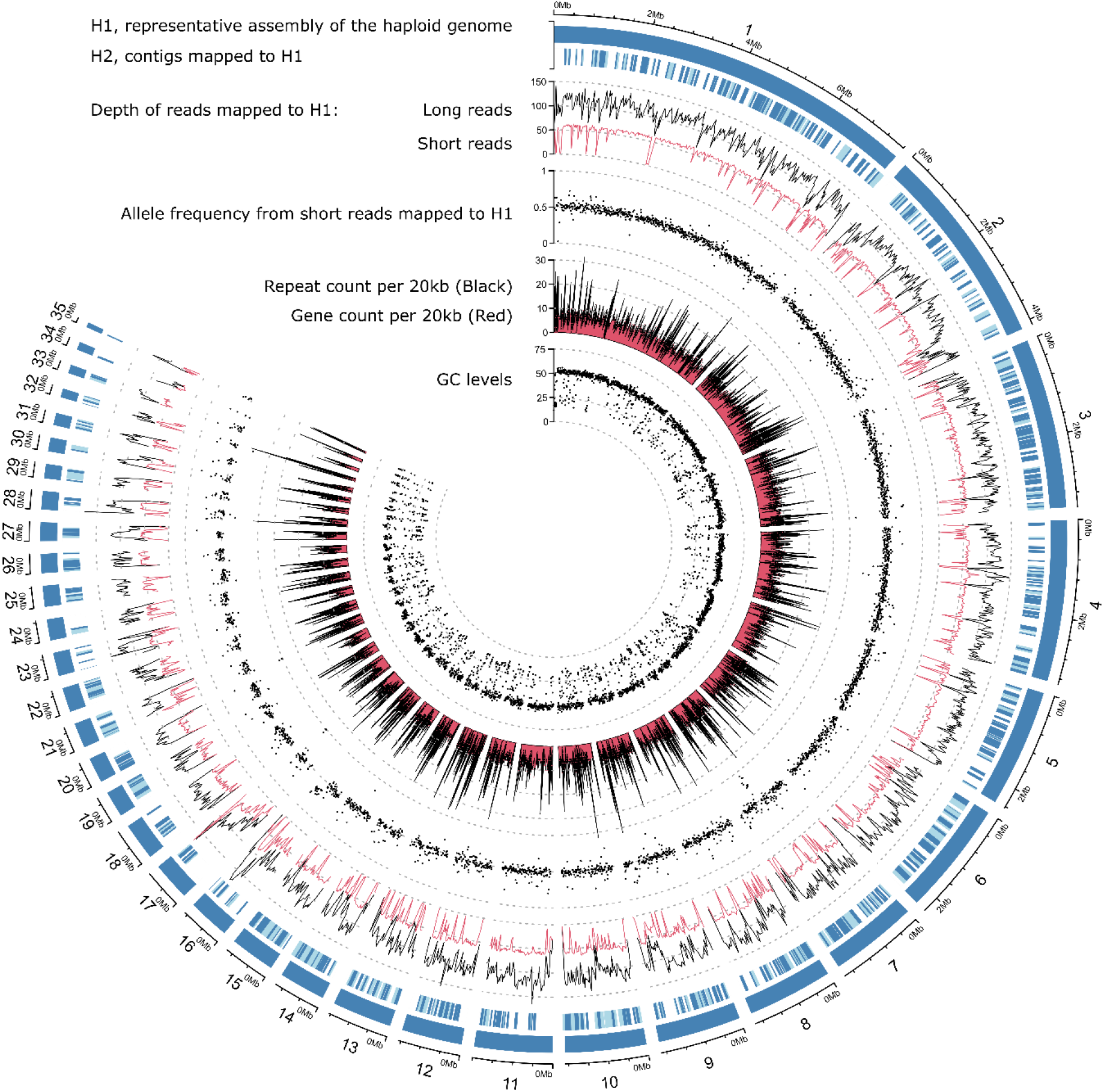
Genome-wide statistics of sequencing data, assembly and traits of the *Metarhizium acridum* strain ARSEF 324 genome. The plot consists of five major tracks; from the outside: i) 549 contigs, 35 contigs placed in the outer band (H1) represent the haploid genome. The remaining contigs (H2) map to H1. H2 contigs are given in light and dark blue, where dark blue indicates the fraction of each contig mapped to H1. The following tracks, show ii) the read depth of long reads (PacBio) and short reads (BGI DBNseq) mapped to H1 (mean depth within 20 kb windows), iii) Allele frequencies calculated from short reads (BGI DBNseq) mapped to H1 (mean within 10 kb windows), iv) Repeat and gene count within 20 kb windows shown in black and red respectively, v) The GC levels within 10 kb windows.

Ideally, the sequencing of an euploid genome should result in an equal read depth across the genome. We mapped both long and short-read data to the H1 assembly, to assess CCNV. The genome wide mean read depth was 54x and 103x (excluding >500x depth, at repetitive elements), for long and short read data respectively (Figure 2). Smaller contigs (<400 kb) tended to have lower read depth, likely as a result poor mapping in repeat dense regions. Contig 34 deviates from the general picture with a mean long-read depth of 52x, the lowest mean depth observed. Contig 34 has a H2 contig that uniquely maps to it, indicating that it is unlikely that contig 34 is a haploid small chromosome. It is possible that it constitutes a haploid segment in the otherwise diploid chromosome. The equal depth across the genome is a strong indication of euploidy across all chromosomes represented in the assembled contigs.

The ploidy of each contig was also assessed by mean ratios of SNV read depth in 10 kb non-overlapping windows. The expectation is that chromosomes with an even ploidy will tend towards a 50:50 distribution across each SNV, while chromosomes with odd ploidy will tend towards a 33:66 or 33:33:33 ratio. All contigs have distribution of ratios with means of 0.5 (50:50) (Figure 2), confirming the diploidy of all contigs. Deviations from 0.5 correlate with read depth of mapped short-read DBNseq data on which the SNV data is based. Mapping of short-reads likewise correlates with the repeat density (Figure 2), because high repeat densities impede the mapping.

The mean GC content across the 44.7 Mb of the H1 contigs is 45% (figure 2, track 5). The GC-content analysed in 100 bp windows shows a bi-modal distribution with peaks at 17.6% and 51.3% (Figure 3A). In the following we delimitate the low and high GC peaks at 33.2%. Repetitive elements explain 81.5% of the low GC peak, 40.5% are unknown repetitive elements, simple repeats 28.7%, retroelements 6.4%, and DNA transposons explain 5.9%. Within low GC windows, 17% were without annotations (Figure 3A). Analysing the dinucleotide composition across the 100 bp windows using the RIP composite index, showed that 33.1% of the genome have a composite index value above zero and are thereby interpreted as affected by RIP. Only 19.3% of the windows with GC content above 33.2% are RIP affected, whereas 95.7% of the low GC peak are affected by RIP (Figure 3B). Across GC levels, 82% of transposable elements are affected by RIP. Similarly, 83% of unknown repeats are RIP affected.

**Fig. 3.**
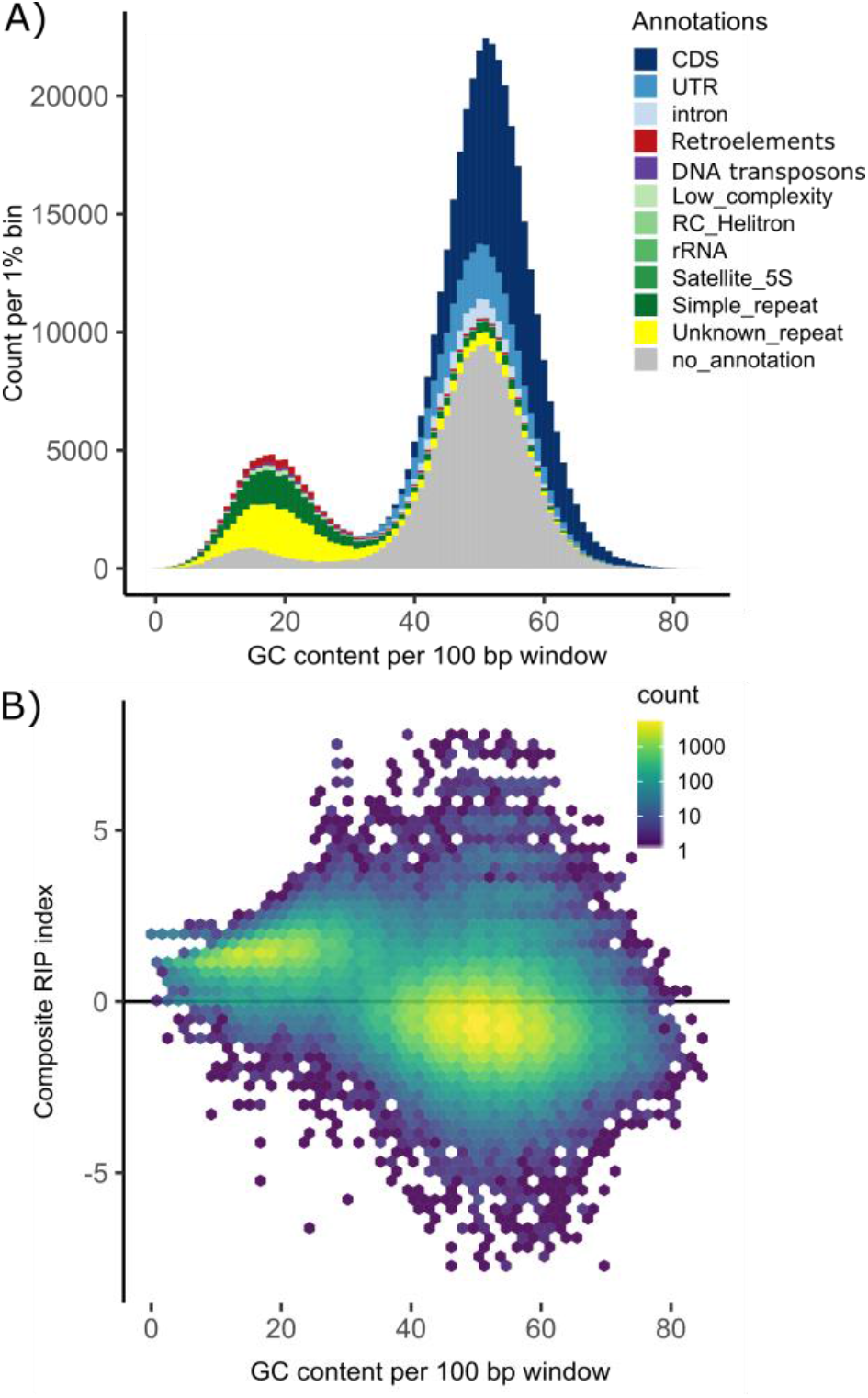
GC content calculated in 100 bp. windows across the 44.7 Mb H1 haploid genome. A) If a window overlapped with more than half of its length with an annotated feature this were signed to the window. The bi-modial distribution have peaks GC-levels of 17.6% and 51.3%. Each peak approximates normal distributions with standard deviations of 5.6 and 6.5, respectively. The fitted normal distributions intersect at 33.2%; this level is set to split high and low GC values. Genomic windows with high GC values is largely comprised of CDS (37.3%), UTR (10.9%), introns (3.3%) and sequence that were without annotations (42.3%). Genomic windows with low GC values is dominated by repetitive elements. Of these repetitive elements 40.5% are unknown, 28.7% are simple repeats, 6.4% are retroelements, 5.9% are DNA transposons. Of the low GC windows 17% were without annotations. B) Composite RIP index as a function of GC content. Composite RIP index >0 are considered RIP affected, equaling 33.1% of the total genome, or 95.7% of the windows within the low GC peak. Combining different indices of RIP will provide a more conservative estimate of the proportion of the genome that is affected by RIP (17.9%, Figure 5).

*M. acridum* ARSEF 324 is homozygous for the MAT1-2 mating-type ideomorph (Figure 4). The two mating-type alleles of *M. acridum* ARSEF 324 and the mating type allele of *M. acridum* CQMa 102 have notable differences in insertions. H1 contains a Mutator-like element (MULE) DNA transposon downstream of MAT1-2-3 not present in H2, contrary H2 harbours a LAGLIDAG homing endonuclease not present in H1. *M. acridum* CQMa 102 contains a WD-repeat domain that is not found in either of the two ARSEF 324 alleles.

**Fig. 4.**
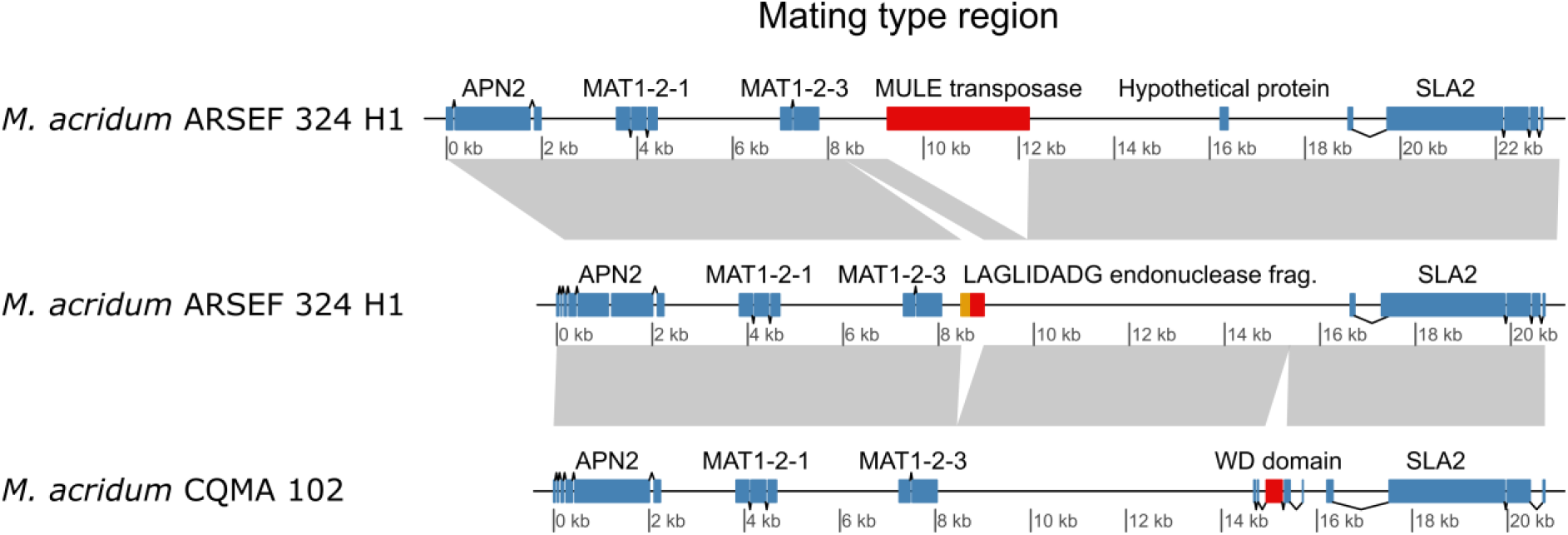
Synteny between mating-type loci of the two alleles *Metarhizium acridum* strain ARSEF 324 and *M. arcridum* strain CQMa 102. The M. acridum strain ARSEF 324 is homozygous for the MAT1-2 mating-type ideomorph. Down-stream of the ARSEF H1 MAT1-2-3 gene, a 3 kb DNA transposon of the ‘MULE’ (Mutator-like elements) superfamily is found, this is not present in ARSEF H2. Mutator transposable elements are among the most mutagenic transposons known (Lisch, 2015; Dupeyron *et al.*, 2019). Conversely, H2 harbours a LAGLIDADG endonuclease fragment (red) together with a fragment of the mitochondrial COX3 gene (orange), not present in H1. Neither of these insertions is present in the CQMa 102 strain, which contains a WD domain not present in ARSEF 324.

### Genome comparison

In order to assess if our haplotig assembly differed from other species within the genus *Metarhizium,* we compared the assembly of strain ARSEF 324 with genomes of from eight *Metarhizium* species (Figure 5). For the phylogenetic analysis, two strains of *Pochonia chlamydosporia* representing the closest known sister taxa to the *Metarhizium* genus was included. The *Metarhizium* genomes presented range in haploid size from 30.8 Mb to 44.7 Mb. The 12.536 predicted genes within ARSEF 324 (H1), fall within the range predicted for other *Metarhizium* isolates, i.e. 8472 to 13,646. A notable difference can be seen in the proportion of repetitive elements between genomes. The median repeat level across the *Metarhizium* isolates is 6.1%, *M. anisopliae* strain JEF-290 has twice this, and the diploid *M. acridum* strain ARSEF 324 has three times this level, with 20.8% of the genome comprised of repetitive elements. The proportion of the genome of *M. acridum* strain ARSEF 324 that is affected by RIP is likewise higher than what was observed for the other genomes analysed (Figure 5). A conservative estimate of the RIP affected region can be given by only including regions that are assessed as RIP affected with all three RIP calculating methods: RIP product, RIP substrate and RIP composite as used above. With this conservative estimate, 17.9% of the *M. acridum* strain ARSEF 324 genome is affected by RIP (calculated in 1000 bp window, in 500 bp steps). Evaluating the genomes ordered in decending order of RIP content, the diploid *M. acridum* strain is followed by *M. acridum* CQMa 102 and *M. rileyi* ARSEF 4871 with 3.8% and 3.6% respectively (Figure 5). The effect of window size on total RIP proportions is shown in Table S3.

**Fig. 5.**
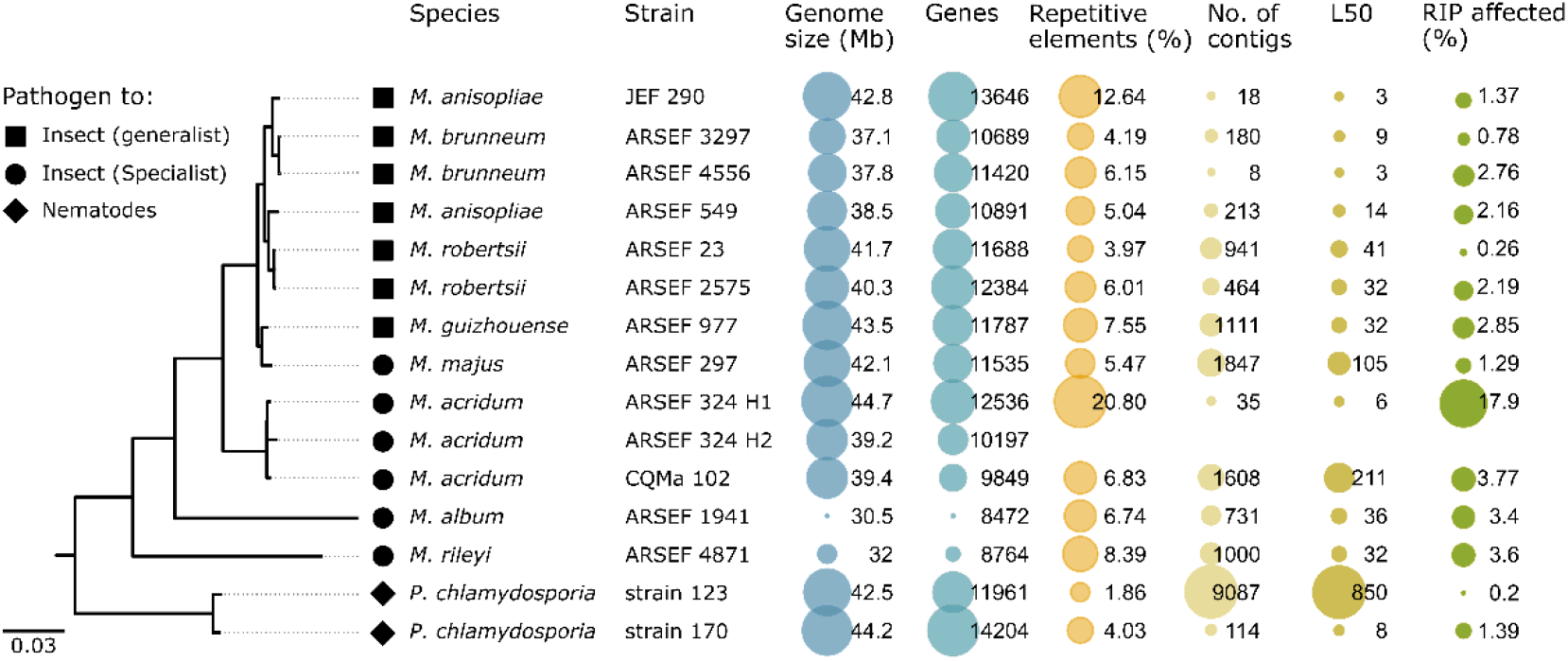
A consensus maximum-likelihood phylogeny based on 444 known single-copy orthologous genes for sequenced *Metarhizium* species and the related *Pochonia chlamydosporia*. All splits within the phylogeny had 100% bootstrap support, except the artificial split between H1 and H2. From the left: ML phylogeny, tip icons indicating the host adaptation, species names and strain id. Genome size and number of genes is retrieved from NCBI Genbank and the protein file available under the accession found in table S2. The proportion of repetitive elements within each isolate was found as described in the method section. The figure highlights that the genome assembly of *M. acridum* ARSEF 324 generated in this study is the longest genome assembly of any *Metarhizium* isolate, and the one with the highest proportion of repetitive elements.

**Fig. 6.**
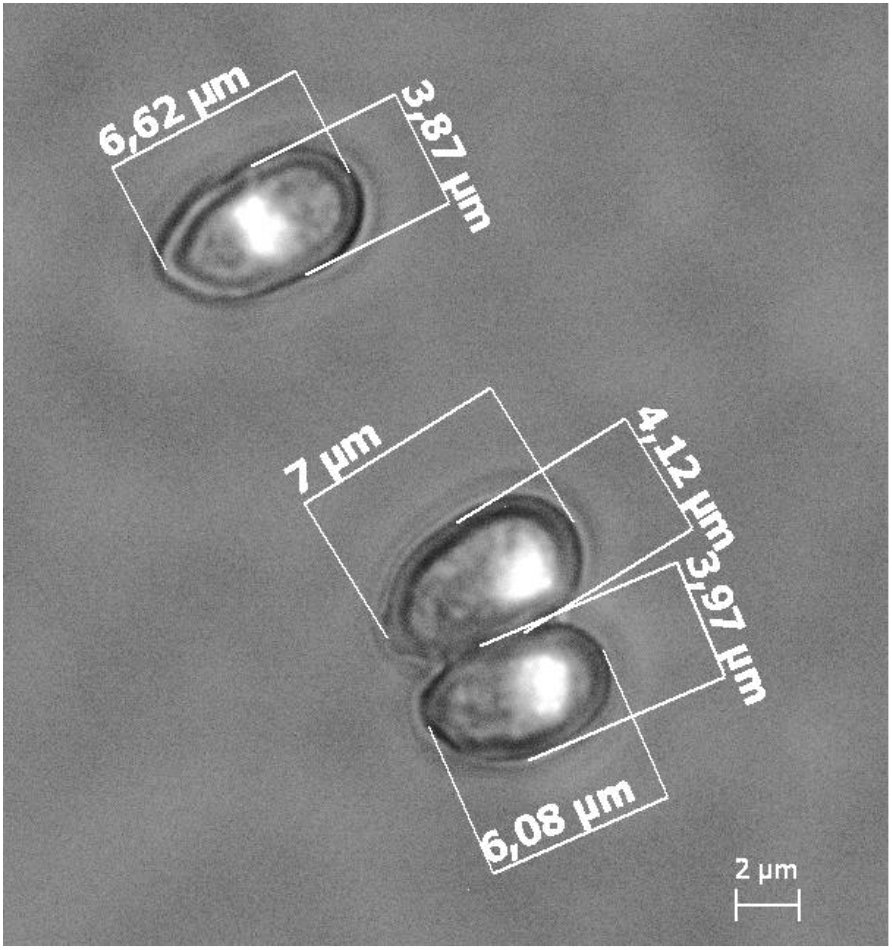
DAPI stained nuclei in *Metarhizium acridum* strain ARSEF 324 conidia. All inspected spores contained monokaryons.

## DISCUSSION

*Metarhizium* species are predominately haploid, with only two taxa within the *M. majus* complex described as stable diploids. Here we present a highly improved genome assembly of *M. acridum* based on long-read sequencing data, which is assembled across repeat regions. This reveals a genome enriched in repetitive elements relative to other *Metarhizium* isolates or species studied. All presented analyses indicate that the sequences *M. acridum* strain ARSEF 324 show the genomic characteristics of a diploid. All 35 contigs of the representative assembled haplotype, were subject to the mapping of smaller contigs and all were heterozygous with allele frequencies between reads around 0.5. The only exceptions are the two minor contigs 17 and contig 34, which could be examples of misassembly, or segmental aneuploidy.

A growing body of evidence indicates that CCNV is a common adaptive trait of many fungi and has been observed at low prevalence in natural populations. The effect on fitness induced by CCNVs has been shown to vary, and can be life-stage specific, and how often these variants are reproductively stable is unknown. We did not see any signs of chromosomal loss to indicate ongoing haploidisation within this diploid strain of *M. acridum*. The absence of chromosome loss suggests that the polyploidisation is recent, or that the strain constitutes a stable diploid. Kepler et al. (2016) found monophyletic clades of diploid isolates in a population genetic study of the *M. majus* species complex (the MGT clade, sensu Bischoff et al 2009). A similar population study could shed light on the prevalence of diploidy or CCNV within *M. acridum*. Alternatively, the diploid stability can be assessed by long-term evolution experiment focusing on possible reversions to the ancestral haploid state. Future research is needed to investigate gene evolution within this strain and identify concerted gene loss levels and rates of pseudo- and neofunctionalisation. If the diploidisation of this strain is not recent, it should be possible to trace divergence between ortholog genes (paralogs, if the diploidisation is the result of endoreplication), where gene duplication did not result in increased fitness and purifying selection.

The highly continuous genome assembly ARSEF 324 includes the highest proportion of repetitive elements of any *Metharhizium* species reported so far. Part of the reason for this is likely to do with the fragmented character of many short read based assemblies and the inherent bias that spring from difficulties in assembling across repeats using these technologies. Our comparative analysis includes, besides the genome assembly of *M. acridum* ARSEF 324, three assemblies based on long-read technologies, i.e. *M. anisopliae* JEF-290 *M. brunneum* ARSEF 4556 and *P. chlamydosporia* 170 (Table S2). Assuming that these assemblies give credible indications of their species’ general repeat content, it is noticeable that two *Metarhizium* species, both belonging to the presumable asexual PARB clade and have considerable less repetitive elements compared with ARSEF 324. This difference between the primarily asexual *M. anisopliae* and *M. brunneum* and the sexual *M. acridum* corroborates the theory that sexual reproduction allows for the proliferation of transposable elements within populations like ‘a sexually-transmitted nuclear parasite’ (Hickey, 1982). Another difference between asexual and sexual fungal species are found in their defense against such selfish DNA elements. The RIP machinery that mutate TE’s, is only active during the sexual cycle and should therefore only be observed in species that sexually recombine and to a less degree in primarily asexually reproducing species. Comparing the repeat content to the proportion of the genome affected by RIP across the *Metarhizium* species show that the two well-assebled genomes of *M. anisopliae* JEF-290 and *M. brunneum* ARSEF 4556 both have a higher ratio of ‘repeats %’ to ‘RIP %’ compared to *M. acridum* ARSEF 324, indicating a lack of an active RIP control of repeats. Both the high repeat content and the active RIP defence supports that *M. acridum* is sexually reproducing, even though no teleomorph is known. The RIP defence of *M. acridum* has shaped the genome by changing the nucleotide composition of approximately one-fifth of the genome, reducing the GC content in affected regions to less than half of the background GC frequency (51%).

The presence of identical mating-type ideomorphs makes it unlikely that the diploidisation results from a mating event as hypothesised of the diploid strains within the *M. majus* species complex. This supports a model where either endoreplication, or allopolyploidsation through parasexuality are responsible for diploidisation. If the former were the case, the duplication event should be old enough for the measurable sequence divergence to have occurred between the homologous genes. The phylogenetic distances between homologous genes within the diploid were comparable to the distance observed between two *M. robertsii* isolates that were isolated with 27 years apart in the USA (Figure S3). This indicates that the observed phylogenetic distance between ARSEF 324 orthologs is consistent with the allopolyploidisation hypothesis. The high prevalence of parasexuality observed in *M. robertsii* renders alloploidisation a likely mode of forming the observed diploid *M. acridum* isolate.

CCNV has thoroughly been linked with both stress and enhanced fitness. It is possible that the high frequency of parasexuality observed in coinfected insects (Riba *et al.*, 1980; Leal-Bertioli *et al.*, 2000; Wang *et al.*, 2011) not just provides a means of recombination but also induce developmental changes that could be beneficial facing a host immune response. If the diploidisation of ARSEF 324 arose from a parasexual fusion, the question remains how stable or long-lived this is. The diploid strain ARSEF 324 is likely to be phenotypically different from haploid *M. acridum* strains. Higher ploidy can enable increased transcription of virulence factors or effectors, and it could be speculated that this could make the strain a more potent biocontrol agent. Diploidy can also work as a shield against detrimental mutation through functional redundancy (Haldane, 1932, p. 110; Orr, 1995) This could explain the strain specific high tolerance to UV-B radiation of ARSEF 324 compared to other *M. acridum* isolates reported by Braga et al. (2001).

While the diploidy reported here for an isolate within *M. acridum* is independent from the cases of diploidy in *M. majus* (Kepler *et al.*, 2016), it is noteworthy that there are several instances of ploidy variation in the genus *Metarhizium.* It is tempting to speculate whether the complex species associations where many isolates are soil-dwelling plant-root endophytes and rhizosphere colonisers as well as potent insect pathogens influence the tendency for ploidy-level variation (St-Leger and Wang 2020). It certainly provides the potential for these fungi to find themselves in stressful microhabitats, and it has been noted that stress is the rule rather than the exemption for *Metarhizium* (Lovett and St. Leger, 2015).

## ACKNOWLEDGEMENTS

HHDFL was supported by a Sapere Aude Grant from the Independent Research Fund Denmark. JES is a CIFAR Fellow in the program Fungal Kingdom: Threats and Opportunities and was supported by NSF DEB 1441715, NSF DEB 1557110, and USDA-NIFA Hatch project CA-R-PPA-5062-H.

## AUTHOR CONTRIBUTIONS

HHDFL initiated the project. MN cultured fungi, performed DNA extraction and initial data QC. KNN performed genome assembly, JFMS and JS carried out genome annotation, KNN and JFMS carried out all other data analyses with input from HHDFL and JS, KNN drafted the manuscript and all authors were involved in the finalisation of the manuscript.

## DATA AVAILABILITY

**Data deposition:** Sequencing data and genome assembly of *Metarhizium acridum* ARSEF 324 has been deposited and are available under the Umbrella Bioproject accession: (Under preparation)

Genbank accession numbers of all genomes used in this study are given in supplementary table S2.

