## Supplementary material for "A NEW CASE OF DIPLOIDY WITHIN A HAPLOID GENUS OF ENTOMOPATHOGENIC FUNGI"

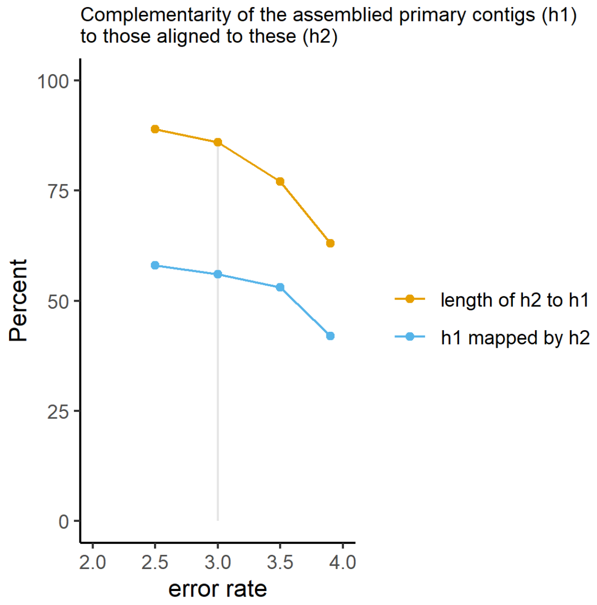


Fig. S1 Complementarity of assembled haplotigs as a function of the Canu assembly parameter ‘correctedErrorRate’.

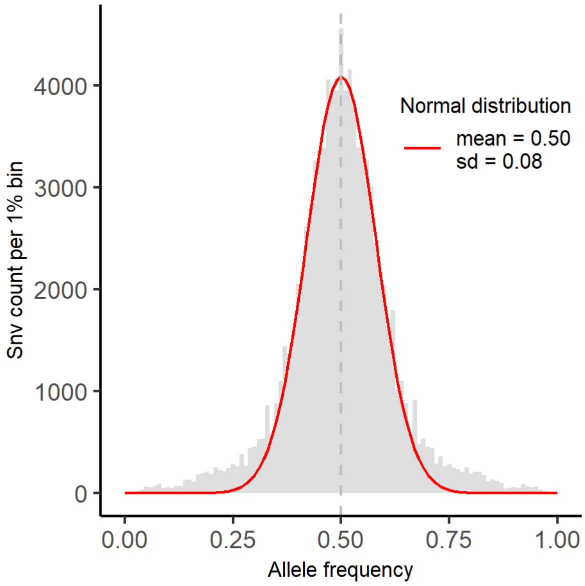


Fig. S2 Allele frequency distribution. The relative read depth of each snv, is given as a frequency, which here has been counted and binned into one per cent bins. A normal distribution has been fitted to the frequency distribution.

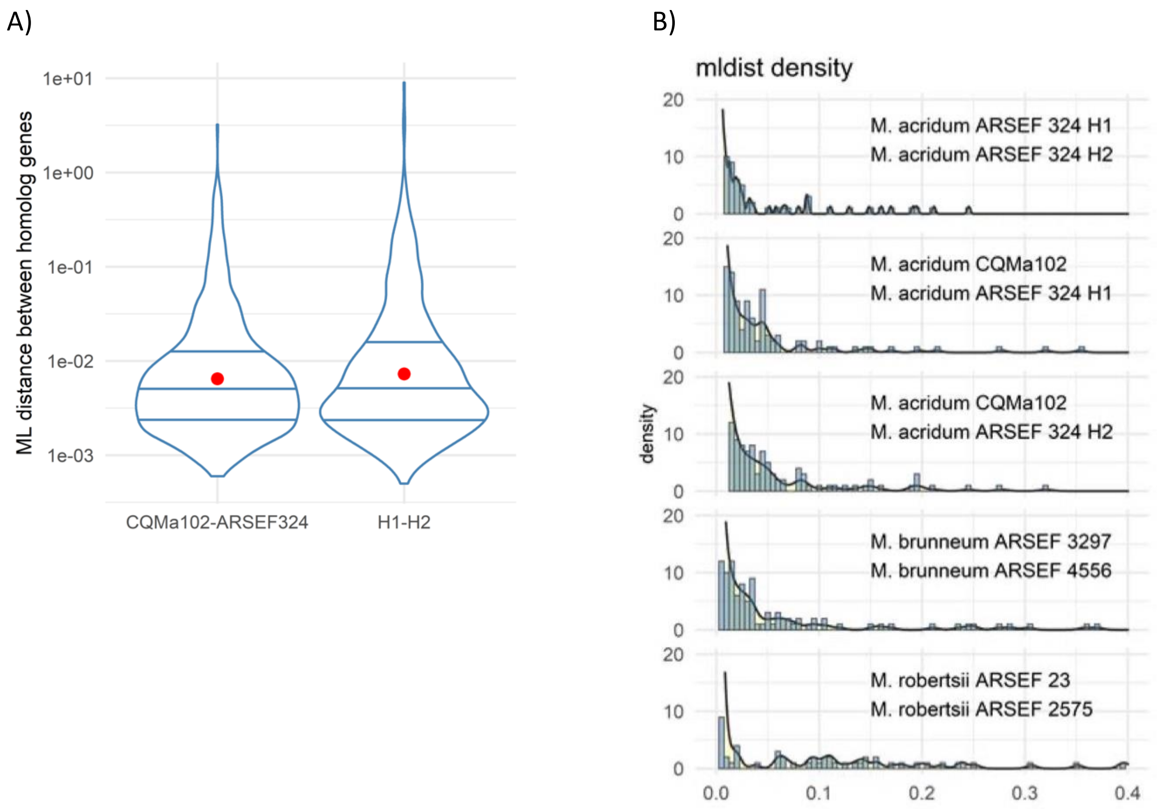


Fig. S3

Divergence between 903 single-copy ortholog gene pairs. Divergence is calculated as the Maximum likelihood (ML) distance between two ortholog genes, given the substitution model fitted based on the multi-sequence alignment. A) Three pairwise comparisons between orthologs within Metarhizium acridum. The comparison between strain CQMa 102 and ARSEF 324 includes distances both to H1 and H2 copies of the same gene. H1 and H2 denote the two copies present in the diploid strain ARSEF 324. The width of the violin plot gives the data density, the horizontal lines, the three quantiles and the red dot the mean ML distance. The y-axis has been log10 transformed. B) Histogram of 238 pairwise ML distances alignment, The superimposed smooth line summarises the density. Maximum-likelihood distances were calculated with IQtree.

Table S1 GenomeScope results: minimums and maximums of estimated values.

| **Property** | **min** | **max** |
|---|---|---|
| Homozygous (aa) | 99.46 % | 99.52 % |
| Heterozygous (ab) | 0.48 % | 0.54 % |
| Genome Haploid Length | 44.97 Mb | 45.04 Mb |
| Genome Repeat Length | 8.17 Mb | 8.18 Mb |
| Genome Unique Length | 36.80 Mb | 36.86 Mb |
| Model Fit | 79.19 % | 96.78 % |
| Read Error Rate | 0.32 % | 0.32 % |

Table S2

| Species | Strain | Haploid genome Size | Protein-coding genes | Repeat Rate (%) | Ploidy | Spore length (mM) | Spore width (mM) | Origin: Host | Origin: Country | Sequencing technology, and genome coverage | GenBank accesstion |
|---|---|---|---|---|---|---|---|---|---|---|---|
| *M. anisopliae* | JEF-290 | 42.8 | 13646 | 12.64 | | | | | | PacBio Sequel, 237x | GCA_013305495.1 |
| *M. anisopliae* | ARSEF 549 | 38.5 | 10891 | 5.04 | | 5.02±0.31 | 2.02±0.18 | | Brazil | Illumina HiSeq, 98x | GCA_000814975.1 |
| *M. robertsii* | ARSEF 2575 | 40.3 | 12384 | 6.01 | | | | Coleoptera: Curculionidae | USA | 454, 25x | GCA_000591435.1 |
| *M. robertsii* | ARSEF 23 | 41.7 | 11688 | 3.97 | | 4.69±0.41 | 2.34±0.20 | Coleoptera: Elateridae | USA | Illumina HiSeq; 454 147x | GCA_000187425.2 |
| *M. brunneum* | ARSEF 4556 | 37.8 | 11420 | 6.15 | | | | Acari: Ixodidae | USA | ONT; Illumina NovaSeq, 100x | GCA_013426205.1 |
| *M. brunneum* | ARSEF 3297 | 37.1 | 10689 | 4.19 | | 4.48±0.26 | 2.09±0.24 | Acari: Ixodidae | Mexico | Illumina HiSeq, 80x | GCA_000814965.1 |
| *M. guizhouense* | ARSEF 977 | 43.5 | 11787 | 7.55 | | 7.19±0.62 | 2.99±0.29 | Coleoptera: Scarabaeidae | France | Illumina HiSeq, 95x | GCA_000814955.1 |
| *M. majus* | ARSEF 297 | 42.1 | 11535 | 5.47 | Diploid | 11.76±1.62 | 2.42±0.34 | Coleoptera: Scarabaeidae | Western Samoa | Illumina HiSeq, 71x | GCA_000814945.1 |
| *M. acridum* | ARSEF 324 H1 | 44.7 | 12536 | 20.80 | Diploid | 6.88±0.64 | 3.75±0.53 | Orthoptera: Acrididae | Australia | PacBio | |
| *M. acridum* | ARSEF 324 H2 | 39.2 | 10197 | | Diploid | 6.88±0.64 | 3.75±0.53 | Orthoptera: Acrididae | Australia | PacBio | |
| *M. acridum* | CQMa 102 | 39.4 | 9849 | 6.83 | | 4.20±0.26 | 2.55±0.22 | Orthoptera: Acrididae | China | Solexa 113x | GCA_000187405.1 |
| *M. album* | ARSEF 1941 | 30.5 | 8472 | 6.74 | | 4.35±0.46 | 1.59±0.26 | Hemiptera: Cicadellidae | Philippines | Illumina HiSeq, 117x | GCA_000804445.1 |
| *M. rileyi* | ARSEF 4871 | 32.0 | 8764 | 8.39 | | | | Host not specified | Korea | Illumina HiSeq, 107x | GCA_001636745.1 |
| *P. chlamydosporia* | 123 | 42.5 | 11961 | 1.86 | | | | | | Illumina HiSeq, 136x | GCA_000411695.2 |
| *P. chlamydosporia* | 170 | 44.2 | 14204 | 4.03 | | | | | | PacBio; Illumina HiSeq, 211x | GCF_001653235.2 |
| Outgroup used for rooting the phylogey, but not included in the visualitation: | | | | | | | | | | | |
| *Epichloe festucae* | Fl1 | | | | | | | | | | GCA_003814445.1 |
| *Villosiclava virens* | UV-8b | | | | | | | | | | GCA_000687475.1 |

Genome features and background informatin of sequenced *Metarhizium* species.

Gray background: values obtained from (Hu *et al.*, 2014) table S1

Table S3

Conservative estimates of the proportion of genomes affected by RIP.

| Isolate<br>Speceis | | Strain | | Genbank accession no. | GC % | RIP % (100bp) | RIP % (200bp) | RIP % (300bp) | RIP % (400bp) | RIP % (500bp) | RIP % (1000bp) | **RIP % (1000bp, 500bp steps)** |
|---|---|---|---|---|---|---|---|---|---|---|---|---|
| M. | anisopliae | JEF | 290 | GCA_013305495.1 | 50.9 | 3.89 | 2.05 | 1.62 | 1.49 | 1.43 | 1.37 | **1.37** |
| M. | anisopliae | ARSEF | 549 | GCA_000814975.1 | 50.95 | 4.58 | | | | | | **2.16** |
| M. | robertsii | ARSEF | 2575 | GCA_000591435.1 | 50.83 | 5.46 | | | | | | **2.19** |
| M. | robertsii | ARSEF | 23 | GCA_000187425.2 | 51.52 | 3.52 | | | | | | **0.26** |
| M. | brunneum | ARSEF | 4556 | GCA_013426205.1 | 50.65 | 5.07 | | | | | | **2.76** |
| M. | brunneum | ARSEF | 3297 | GCA_000814965.1 | 51.52 | 4.02 | | | | 0.97 | 0.83 | **0.78** |
| M. | guizhouense | ARSEF | 977 | GCA_000814955.1 | 49.65 | 4.49 | | | | | | **2.85** |
| M. | majus | ARSEF | 297 | GCA_000814945.1 | 50.98 | 4.74 | | | | | | **1.29** |
| M. | acridum | ARSEF | 324 | H1 | 45.5 | 19.57 | 18.2 | 17.94 | 17.92 | 17.92 | 17.93 | **17.9** |
| M. | acridum | ARSEF | 324 | H2 | 43.6 | 22.16 | | | | | | **20.9** |
| M. | acridum | CQMa | 102 | GCA_000187405.1 | 49.81 | 5.46 | 4.32 | 4.15 | 4.12 | 4.08 | 3.9 | **3.77** |
| M. | album | ARSEF | 1941 | GCA_000804445.1 | 52.8 | 4.39 | 3.23 | 3.06 | 3.1 | 3.18 | 3.43 | **3.4** |
| M. | rileyi | ARSEF | 4871 | GCA_001636745.1 | 49.92 | 4.48 | | | | | | **3.6** |
| Pochonia | chlamydosporia | strain | 123 | GCA_000411695.2 | 49.87 | 2.94 | | | | | | **0.2** |
| Pochonia | chlamydosporia | strain | 170 | GCA_001653235.2 | 49.52 | 4.19 | | | | | | **1.39** |

The proportion assigned as RIP affected, were so done based on three RIP indices calculated with the web-tool: http://theripper.hawk.rocks).

All analyses were made with non-overlapping window, i.e same window size and step size. Except for RIP % (1000bp, 500bp steps), which is the default setting.
